# *Drosophila melanogaster* pigmentation demonstrates adaptive phenotypic parallelism over multiple timescales

**DOI:** 10.1101/2024.08.09.607378

**Authors:** Skyler Berardi, Jessica A. Rhodes, Mary Catherine Berner, Sharon I. Greenblum, Mark C. Bitter, Emily L. Behrman, Nicolas J. Betancourt, Alan O. Bergland, Dmitri A. Petrov, Subhash Rajpurohit, Paul Schmidt

## Abstract

Populations are capable of responding to environmental change over ecological timescales via adaptive tracking. However, the translation from patterns of allele frequency change to rapid adaptation of complex traits remains unresolved. We used abdominal pigmentation in *Drosophila melanogaster* as a model phenotype to address the nature, genetic architecture, and repeatability of rapid adaptation in the field. We show that *D. melanogaster* pigmentation evolves as a highly parallel and deterministic response to shared environmental gradients across latitude and season in natural North American populations. We then experimentally evolved replicate, genetically diverse fly populations in field mesocosms to remove any confounding effects of demography and/or cryptic structure that may drive patterns in wild populations; we show that pigmentation rapidly responds, in parallel, in fewer than fifteen generations. Thus, pigmentation evolves concordantly in response to spatial and temporal climatic gradients. We next examined whether phenotypic differentiation was associated with allele frequency change at loci with established links to genetic variance in pigmentation in natural populations. We found that across all spatial and temporal scales, phenotypic patterns were associated with variation at pigmentation-related loci, and the sets of genes we identified in each context were largely nonoverlapping. Therefore, our findings suggest that parallel phenotypic evolution is associated with distinct components of the polygenic architecture shifting across each environmental gradient to produce redundant adaptive patterns.

## Introduction

Characterizing the dynamics of adaptation in the field is a central goal of evolutionary biology. To better understand how populations may respond to environmental change, it is necessary to define the tempo and repeatability of evolution at both the phenotypic and genomic level. These questions have been explored across multiple scales, from macroevolutionary patterns and broad adaptive radiations (Knoll & Carroll, 1999) to controlled laboratory experiments detailing microbial evolution (Lenski et al., 1991). Examining geographic clines in phenotypic and genotypic variation represents a classic approach to studying adaptation (Endler, 1977), and clines are often viewed through a macroevolutionary lens (Allen, 1877). Yet, both the exact timescale over which local adaptation occurs, and the respective roles of selection and demography in maintaining clines, are often obscured. Therefore, to understand how populations respond to the environment over ecological timescales, recent studies have leveraged high resolution temporal sampling as a powerful approach to decode genetic changes that may contribute to adaptive evolution in the field (Barrett et al., 2008; Barrett et al., 2019; Bitter et al., 2024; Enbody et al., 2023; Jones et al., 2018; Nosil et al., 2018; Roberts Kingman et al., 2021; Rudman et al., 2022). However, rarely have these approaches been combined to examine adaptation across multiple scales, and integrating spatial and temporal patterns provides an opportunity to address several fundamental questions about the dynamics of adaptation in natural populations. In particular, we can test whether patterns of adaptive change are parallel across multiple environmental gradients that arise over space and time, define the timescales over which adaptive patterns are established, and examine how allele frequencies for putatively causal loci covary with phenotypic patterns. Therefore, to explore the pace and predictability of adaptation in the field and integrate phenotypic patterns with genomic shifts, we studied the evolution of a complex trait across multiple timescales in *D. melanogaster*.

*Drosophila* species have been a classic model for exploring evolution over ecological timescales (e.g., Dobzhansky, 1943). *D. melanogaster* exhibits signatures of rapid evolution across a broad array of fitness-related phenotypes (Behrman et al., 2015; Behrman et al., 2018; Erickson et al., 2020; Rajpurohit et al., 2017; Rajpurohit et al., 2018; Schmidt & Conde, 2006). Additionally, seasonally cycling polymorphisms have been detected in populations of *D. melanogaster*, representing genomic signatures of rapid evolution (Bergland et al., 2014; Bitter et al., 2024; Machado et al., 2021; Nunez et al., 2024; Rudman et al., 2022), and studies have begun to integrate rapid phenotypic adaptation with underlying patterns of genetic variation in this system (Erickson et al., 2020; Rajpurohit et al., 2018). We thus aimed to probe for adaptive parallelism across multiple scales using a complex, fitness-related trait with a well-characterized genetic basis. Thousands of polymorphisms vary latitudinally and seasonally in *D. melanogaster* (Bergland et al., 2014; Machado et al., 2021; Rudman et al., 2022), so studying a trait with a known genetic basis would enable our analyses to target variation with established links to phenotype. We selected *D. melanogaster* pigmentation as our model trait, and our motivation for studying pigmentation was threefold: it exhibits extensive phenotypic variation in wild populations (Kronforst et al., 2012), it is putatively adaptive (Bastide et al., 2014; Pool & Aquadro, 2007), and it has a well-studied genetic architecture (Wittkopp et al., 2003).

*Drosophila* pigmentation varies dramatically in natural environments both across and within species (Kronforst et al., 2012). Latitudinal clines for *D. melanogaster* pigmentation have been described in Europe, India, and Australia, in which melanization increases with latitude (Das, 1995; David et al., 1985; Munjal et al., 1997; Parkash et al., 2008; Rajpurohit et al., 2008a; Telonis-Scott et al., 2011). Additionally, pigmentation has been shown to increase with altitude in Sub-Saharan Africa and India (Bastide et al., 2014; Dev et al., 2013; Munjal et al., 1997; Parkash et al., 2008; Pool & Aquadro, 2007; Rajpurohit et al., 2008a). These repeated clinal patterns suggest pigmentation is likely an adaptive response to environmental conditions that vary predictably over latitudinal and altitudinal gradients, such as temperature (Freoa et al., 2023; Munjal et al., 1997). Crucially, the pathways underlying melanin and sclerotin biosynthesis in each abdominal tergite are well-characterized (Kopp et al., 2003; True, 2003; Wittkopp et al., 2003). Melanization is influenced by major-effect genes including *tan, ebony, yellow, bab1*, and *bab2* (Couderc et al., 2002; Kopp et al., 2000; True et al., 2005; Wittkopp et al., 2002; Wright, 1987), and natural variation in pigmentation has been mapped to single nucleotide polymorphisms (SNPs) in both major- and minor-effect loci (Bastide et al., 2013; Dembeck et al., 2015; Endler et al., 2016). Therefore, given that segregating variants for canonical pigmentation genes exist in natural populations of *D. melanogaster*, we hypothesized that adaptive shifts in pigmentation may be driven by repeated selection at common variants of major-effect loci, similar to alleles of *Mc1r* and *Agouti* driving adaptive pigmentation patterns in wild populations of mice (e.g., Barrett et al., 2019; Hoekstra et al., 2006; Manceau et al., 2011). However, the canonical *Drosophila* pigmentation genes are also highly pleiotropic (Massey & Wittkopp, 2016; Wittkopp & Beldade, 2009), which may constrain selection over ecological timescales (Paaby & Rockman, 2013). In this case, pigmentation patterns may not be driven by shifts in major-effect pigmentation genes, but instead by selection across variants that are less subject to pleiotropic constraints (Kostyun et al., 2019).

We studied *D. melanogaster* pigmentation in wild populations to test several fundamental hypotheses about the dynamics of adaptation. We sought to characterize whether patterns of phenotypic and genomic evolution are parallel across multiple scales, and to identify whether timescale modulates how patterns of genetic variation map to phenotype. We first examined whether pigmentation patterns are concordant across three environmental gradients with shared conditions, as would be expected if this trait adapts predictably to drivers such as temperature. After establishing parallelism in the phenotypic response across multiple spatiotemporal scales, we then explored whether shifts in known, major- or minor-effect pigmentation genes consistently underpin observed phenotypic patterns. This enabled us to determine whether repeated phenotypic adaptation is associated with variation in canonical pigmentation genes across each spatial and temporal scale examined.

## Methods

### Fly collection and maintenance

#### Wild orchard populations

*D. melanogaster* individuals were collected from orchard populations located along the east coast (U.S.A.) by either direct aspiration or bait-and-sweep net methods. Gravid females were sorted into isofemale lines in the field; once resulting progeny eclosed, lines were typed to species. Latitudinal populations were sampled in the spring across six sites: Lancaster, MA (42.45N, 71.87W); Media, PA (39.88N, 76.30W); Charlottesville, VA (38.02N, 77.47W); Athens, GA (33.95N, 83.66W); Jacksonville, FL (30.20N, 81.66W); and Homestead, FL (25.28N, 80.45W). Seasonal collections were done in Media, PA, over six years (2010-2015). Early-season collections were made over the first week of June, and late-season collections were completed in early November (Behrman et al., 2015; Behrman & Schmidt, 2022; Rajpurohit & Schmidt, 2016). All populations were maintained long-term at 24°C, 12L:12D photoperiod, on standard cornmeal-molasses-agar medium.

#### Experimental orchard populations

On June 30, 2016, we founded 10 replicate, outdoor mesocosms with 500 males and 500 females from an outbred founder population that was established by recombining a panel of 80 inbred lines, as described by Rudman et al. (2022). Each 8m^3^ mesocosm contains a dwarf peach tree and vegetative ground cover to provide a semi-natural environment for fly rearing. The experiment was conducted from June 30-October 5, 2016, and each population was provided 1 liter of cornmeal-molasses medium each week. To assess evolutionary patterns, *D. melanogaster* eggs were sampled from each mesocosm at three timepoints (August 5, September 5, and October 5). DNA was sequenced at all three timepoints, and flies were preserved for pigmentation measurements at two timepoints (Aug. 5 and Oct. 5). 100 females were pooled from each mesocosm to conduct whole genome sequencing (using the DNA extraction, sequencing, and analysis pipelines described in Rudman et al. (2022) and Bitter et al. (2024)). To measure melanization at each timepoint, flies sampled from each mesocosm were reared for 2 generations in laboratory, common garden conditions (25°C, 12L:12D, ∼30 eggs per vial), to control for any effects of developmental plasticity. Five-day old F3 individuals corresponding to each mesocosm and timepoint were stored in 80% EtOH at -80°C for later phenotyping.

### Scoring fly samples for abdominal pigmentation

#### Wild orchard populations

We scored female abdominal pigmentation following common garden treatment using a lateral view of the abdomen, giving values from 0 (0% of the tergite is melanized) to 10 (100% melanized) for each of seven abdominal tergites as described by David et al. (1990) (Fig. 1). We only scored female flies, as abdominal pigmentation in male flies is less variable among individuals. To measure latitudinal patterns, we selected a total of 17-30 isofemale lines from each population, and we scored ten individual females per line. To measure seasonal patterns, we sampled 10-20 isofemale lines from Media, PA, at early- and late-season timepoints, and scored 5-10 individuals per line. We then calculated the mean pigmentation score for each isofemale line across seasons and years. Latitudinal pigmentation patterns were analyzed in R (v.4.4.0) with a linear mixed effects model using the package ‘nlme’ (v.3.1.164). Here, latitude was modeled as a fixed effect and isofemale line was modeled as a random effect, according to the formula: lme(Pigmentation Scores ∼ Latitude, random=∼1|Isofemale Line). Similarly for the seasonal (Media, PA) populations, we examined pigmentation patterns with a linear mixed effects model where year, season (early or late), and their interaction were modeled as fixed effects, and isofemale line was included as a random effect: lme(Pigmentation Scores ∼ Year*Season, random=∼1|Isofemale Line).

**Fig 1.**
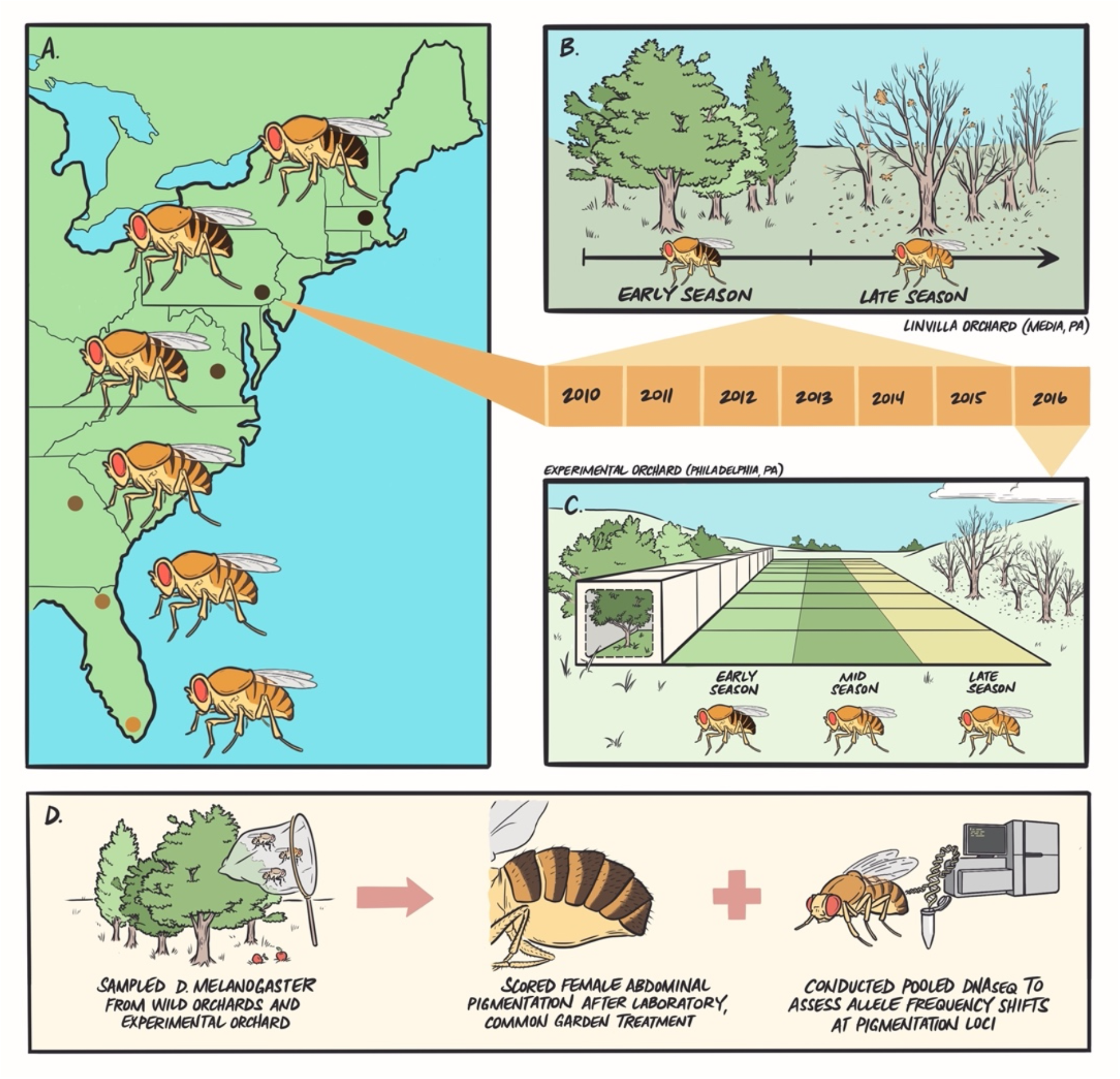
Experimental overview. (A) We sampled flies from six wild orchard populations ranging from Homestead, FL, to Lancaster, MA, and established isofemale lines in the laboratory. (B) We returned to a focal orchard in Media, PA, at early- and late-season timepoints and collected flies to capture evolutionary patterns following winter and summer conditions. (C) We then seeded outdoor mesocosms (N=9) with an outbred population originating from early-season collections in Media, PA, and sampled flies at the end of summer (mid-season) and fall (late-season) to determine if seasonal patterns are recapitulated in experimental populations controlled for migration, drift, and cryptic population structure. (D) Across each wild or experimental context, we sampled flies, established lines in the lab, completed common garden treatment to remove environmental effects, and scored females for abdominal pigmentation. We also conducted pooled DNA sequencing on additional flies sampled from each population to map genomic patterns for candidate pigmentation SNPs.

#### Experimental orchard populations

Abdominal pigmentation of *D. melanogaster* females was scored using the rubric described above (David et al., 1990). The three most distal tergites were scored (#5-7), as they show the highest degree of variation. We first scored 4 replicates of 20 females sampled from our ancestral founder population. For the August 5 and October 5 field-evolved timepoints, 20 individuals from each mesocosm were randomly selected and scored from F3 samples stored at -80°C; we then calculated a mean pigmentation score for each mesocosm. One of our initial 10 mesocosms was lost (“S2”), so we scored patterns for the remaining 9 mesocosms. We ran a linear mixed effects model in R (‘nlme’, v.3.1.164) for pigmentation scores in each mesocosm between the summer and fall timepoints. Here, timepoint was included as a fixed effect and was modeled as an ordered categorical variable, and mesocosm was included as a random effect nested within timepoint. We used the formula: lme(Pigmentation Scores ∼ Timepoint, random=∼1|Mesocosm/Timepoint). We included scores from our mid-(Aug. 5) and late-season (Oct. 5) timepoints in this model, but we excluded the founder population because we did not score pigmentation for the subsetted founder flies that seeded each cage.

### Genomic Analyses of Sequencing Data

#### Selection of candidate SNPs

We based our candidate SNP selection on naturally segregating variants identified by two previous GWAS of abdominal pigmentation (Bastide et al. 2013; Dembeck et al. 2015). Bastide et al. (2013) collected wild samples in Austria and Italy, and Dembeck et al. (2015) used inbred lines of the Drosophila Genetic Reference Panel (DGRP). These GWAS focused on different tergites: Bastide et al. (2013) looked at pigmentation of the 7th tergite, and Dembeck et al. (2015) focused on the 5th and 6th and their relationship. Since our pigmentation scoring considered the 5th, 6th, and 7th tergites, we included all significant SNPs from both studies (a total of 249 SNPs associated with 111 genes). We based our SNP coordinates on the *D. melanogaster* Release 5 (dm3) coordinate system, which required us to convert SNP coordinates in the latitudinal and seasonal genomic datasets from Release 6 back to Release 5. From the initial paper describing genome Release 6 (dos Santos et al., 2015), we downloaded the associated FlyBase file (http://flybase.org/reports/FBrf0225389.htm) that maps Release 5 to Release 6 and wrote a pipeline to convert SNP coordinates between releases.

#### Calculating a metric of latitudinal clinality of candidate SNPs

To measure the degree of clinality for each SNP, we performed a linear regression (‘lm’ in R) on SNPs’ frequencies across latitudes. Our metric for clinality was the absolute value of the regression coefficient of the resulting model. Latitudinal samples we analyzed were collected from Homestead, FL (2010), Atlanta, GA (2014), Charlottesville, VA (2014), Media, PA (2014) and Lancaster, MA (2014); however, we did not have genomic data available for Jacksonville, FL, to include in our analyses. Data for these samples was sourced from the DEST dataset (Kapun et al., 2021), and corresponding sample IDs were: “FL_ho_10_spring”, “GA_at_14_spring”, “VA_ch_14_spring”, “PA_li_14_spring”, and “MA_la_14_spring.” We excluded any SNPs that were present in less than 3 of the 5 samples.

#### Calculating a metric of seasonality of candidate SNPs in wild orchard populations

We determined that the allele frequency change from spring to fall and the repeatability of this shift were the key components of seasonality. Therefore, we considered periods where both a spring and a fall sample existed in a given year, and we calculated the magnitude of allele frequency change between seasons. We then averaged across these spring-fall transitions to get the yearly seasonal change. Seasonal samples were collected in Media, PA, from 2009-2015 (DEST dataset, Kapun et al., 2021), and corresponding sample IDs were: “PA_li_09_spring”, “PA_li_09_fall”, “PA_li_10_spring”, “PA_li_10_fall”, “PA_li_11_spring”, “PA_li_11_fall”, “PA_li_12_spring”, “PA_li_12_fall”, “PA_li_14_spring”, “PA_li_14_fall”, “PA_li_15_spring”, and “PA_li_15_fall.” Notably, we did not measure pigmentation patterns during the 2009 season, so this year is only represented in our genomic analyses; additionally, our genomic analyses did not include samples from 2013, which were included in our phenotypic analyses. We excluded SNPs with data in fewer than 3 out of a possible 6 spring-fall pairs.

#### Calculating a metric of seasonality of candidate SNPs in outdoor mesocosm populations

For experimental seasonal measurements, we performed a linear regression of allele frequencies across the three timepoints collected during 2016. While we only had pigmentation measurements available from August 5 and October 5 to include in our phenotypic analyses, DNA was sequenced from an additional timepoint (September 5), which was included in our genomic analysis. We designated the absolute value of the regression coefficient as the metric for seasonal change to identify SNPs that shifted in a large, concordant direction over time. We excluded SNPs with missing data in any of the samples, as there were only 3 timepoints.

#### Selection of matched SNPs

We selected a group of at least 200 matched SNPs for each candidate SNP to establish a null model of expected clinal and seasonal patterns. Matched SNPs were: located on the same chromosome further than 50KB from the candidate SNP, found in the same type of region (exon, 5’UTR/3’UTR, intron, gene, or intergenic, in order of priority), and had the same relationship to the 6 major inversions that occur in wild North American populations (within an inversion, within 500KB of a breakpoint, or further than 500KB). After matching on the above characteristics, SNPs with an average frequency across samples within 0.01 of the average frequency of the candidate SNP were chosen as the matched SNP pool for that candidate SNP. If fewer than 200 SNPs were in the pool for a candidate SNP, we increased the possible difference in frequency until the pool reached at least 200 SNPs. However, no matched SNP had a frequency difference of more than 0.07 from its candidate SNP.

#### Tests of overall clinality and seasonality of the entire set of candidate SNPs

To determine the clinality and seasonality of candidate SNP frequency changes as a group, we leveraged matched SNPs to create 1,000 bootstrap-resampled groups equal in size to the number of candidate SNPs that passed the minimum threshold for each metric. Each matched group included one matched SNP drawn randomly from the pool of each candidate SNP with sufficient data. The averages of the clinal or seasonal metrics within each of the 1,000 matched groups provided a null distribution of 1,000 values, which was used to calculate the *p*-value of the average of the candidate group. By creating these bootstrap resamples that matched the properties of the original candidate SNPs, we determined if the candidate SNPs as a group were more likely to follow clinal and seasonal patterns observed in the pigmentation phenotype, given enrichment for a significant magnitude of absolute change.

#### Tests of clinality and seasonality of individual candidate SNPs

To test if each individual SNP varied more latitudinally or seasonally than expected, we drew a sample of 100 matched SNPs from its pool, and we repeated this 100 times. For each selection of 100 matched SNPs, we determined the proportion of matched SNPs with a metric greater than the candidate SNP. We considered a SNP significant if it was within the top 10% of matched SNPs in at least 90 trials and within the top 5% in at least half of the 100 trials, providing a significance cutoff of α=0.05 ± 0.05.

#### Data analysis and figures

All phenotypic and genomic analyses were conducted in R (v.4.4.0) using the packages tidyverse (v.2.2.0), reshape2 (v.1.1.4), readxl (v.1.4.3), and nlme (v.3.1.164). Figures were made using Keynote (v.8.1) and R (v.4.4.0) using the packages ggplot2 (v.3.5.1), emmeans (v.1.10.2), and plotrix (v.3.8.4). Fig. 1 was illustrated by Dr. Rush Dhillon (https://www.rushstudio.ca/).

## Results

### Abdominal pigmentation increases with latitude in North America

We first determined whether *D. melanogaster* pigmentation varies latitudinally in North America, a previously uncharacterized transect. We predicted melanization would increase with latitude, consistent with documented latitudinal clines for abdominal pigmentation in *D. melanogaster* on other continents (Das, 1995; Munjal et al., 1997; Parkash et al., 2008; Rajpurohit et al., 2008a; Telonis-Scott et al., 2011). We measured female abdominal pigmentation for six populations along the East Coast of the U.S., and all populations were phenotyped under laboratory common garden conditions to identify shifts in pigmentation driven by genetic changes (see Methods; Fig. 1A,D). Our results aligned with previously established clines: melanization increased positively with latitude in North America (*F*_5,157_ = 11.17, *p* < .0001; Fig. 2A,D; Table S1A). These parallel and independent clines across continents suggest pigmentation evolves adaptively in response to environmental conditions that vary predictably with latitude (Endler, 1977). However, both demography and local adaptation can drive clinal patterns, and allele frequency clines in *D. melanogaster* appear driven by local adaptation as well as admixture (Bergland et al., 2016). Therefore, we next asked whether pigmentation evolves in response to seasonal change in a focal orchard, and whether seasonal and latitudinal patterns are concordant based on shared putative environmental drivers.

**Fig 2.**
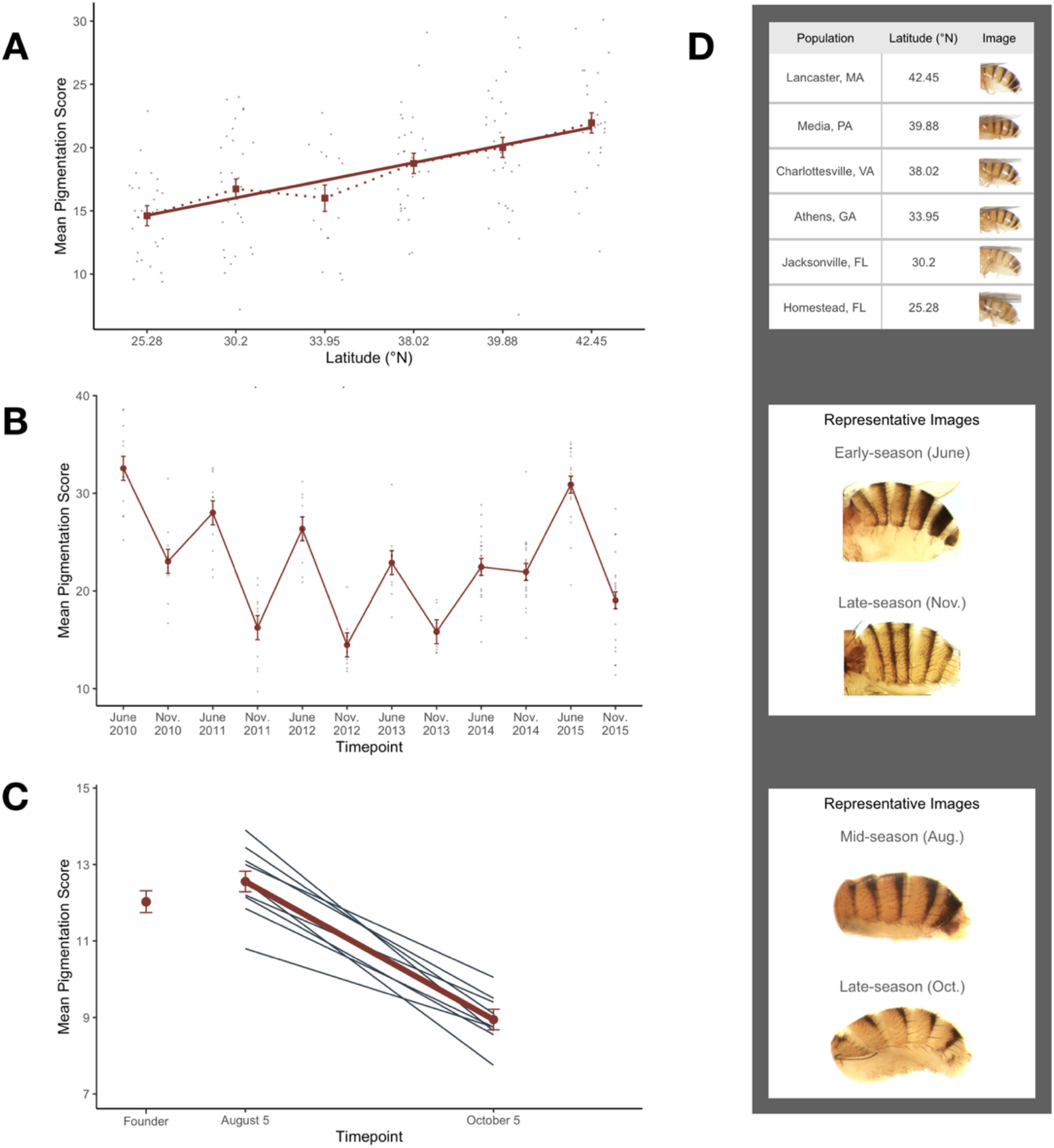
Phenotypic patterns and representative images for *D. melanogaster* populations in each natural context. (A) *D. melanogaster* pigmentation increases significantly with latitude. (B) A population of flies sampled from Media, PA, exhibit predictable and significant decreases in melanization from June to November over six years. (C) Decreases in pigmentation over seasonal time are recapitulated in semi-natural mesocosms (N=9) located in Philadelphia, PA, and female flies adapt to exhibit significantly lighter pigmentation between August 5^th^ and October 5^th^. The average pigmentation score of our spring-derived ancestral population (“Founder”) is also pictured. In all plots, mean pigmentation score and standard error are depicted alongside either points representing isofemale line means (A,B) or individual mesocosm means (C). (D) Representative images corresponding to each population (from top to bottom): across latitudes, across seasons in wild populations, and across seasons in experimental populations.

### Pigmentation varies predictably over seasonal timescales in a wild orchard population

We explored the temporal dynamics of melanization by examining flies collected from Media, Pennsylvania, across seasons in six successive years. Here, we tested the hypothesis that pigmentation evolves in response to conditions, such as temperature, that vary in parallel across latitude and season. Additionally, we aimed to establish the timescale over which pigmentation responds to environmental gradients. We collected flies in early June and early November from 2010-2015 and scored female abdominal pigmentation following common garden treatment (Fig. 1B,D). If pigmentation evolves in response to climate-associated environmental factors, we hypothesized that early-season flies (“winter-evolved”) would have darker pigmentation similar to flies from higher latitudes. Likewise, we predicted that late-season flies (“summer-evolved”) would exhibit lighter pigmentation in parallel to populations originating from lower latitudes. However, the evolution of pigmentation could be constrained over short timescales, or the environmental pressures driving variation in pigmentation may not be shared across spatial and temporal axes. In this case, we would expect to see seasonal patterns that are distinct from latitudinal patterns.

We found that *D. melanogaster* females evolve seasonally and become significantly less melanized in the fall relative to the spring (*F*_1,148_ = 174.43, *p* < .0001; Fig. 2B,D; Table S1B), and this seasonal pattern was reset and repeated across the six years recorded. The magnitude of pigmentation change varied with year (*F*_5,148_ = 11.52, *p* < .0001), with 2014 showing a dampened response. Thus, abdominal pigmentation evolves predictably and in parallel over the ∼15 generations that span spring to fall in this species (Behrman & Schmidt, 2022). The directionality of seasonal change is consistent with latitudinal patterns, supporting the hypothesis that environmental factors that vary in parallel over space and time (e.g., temperature) may be the primary drivers of phenotypic shifts. While this provided further evidence that pigmentation evolves adaptively in response to environmental heterogeneity, we still could not rule out effects of migration or cryptic population structure. Therefore, we utilized experimental evolution in field mesocosms to eliminate these confounding factors and directly examine the adaptive dynamics of pigmentation over ecological timescales.

### Replicate experimental populations of D. melanogaster exhibit highly rapid and parallel adaptation of pigmentation in field mesocosms

We characterized seasonal patterns of abdominal pigmentation in replicate experimental populations located in Philadelphia, Pennsylvania. The populations were housed in outdoor mesocosms to isolate the role of adaptation in driving phenotypic evolution, while maintaining a semi-natural environment (Grainger et al., 2021; Rajpurohit et al., 2017; Rajpurohit et al., 2018; Rudman et al., 2019; Rudman et al., 2022). We seeded replicate mesocosms with an outbred founder population derived from spring collections in our focal orchard (Media, PA), ensuring our starting population had a similar genetic background to local wild populations (see Methods; Fig. 1C). We then measured seasonal evolution by collecting individuals from each mesocosm at each timepoint and scoring female pigmentation after laboratory, common garden treatment (Fig. 1D). If the variance in pigmentation among latitudinal and seasonal populations is driven by selection, we anticipated that we would observe the same pattern we did in wild populations: melanization would decrease from spring to fall in the experimental orchard, similar to wild-caught Pennsylvania flies. In accordance with our prediction, we found late-season flies were indeed less melanized relative to early-season flies across replicate mesocosms (*F*_1,8_ = 111.26, *p* < .0001; Fig. 2C,D; Table S1C), showing a concordant magnitude and directionality of phenotypic change to the wild populations. These parallel phenotypic patterns were observed across nine independent, replicate populations, strongly supporting our hypothesis that the observed shifts in melanization are adaptive.

Together, our findings indicate that pigmentation exhibits a strikingly parallel phenotypic response to environmental conditions that vary across all timescales examined, and this represents the first demonstration that *D. melanogaster* pigmentation adapts rapidly in response to seasonally changing environments. Defining these adaptive patterns set the stage for us to characterize the underlying genomic architecture in each spatiotemporal context. Therefore, we next studied evolutionary patterns at the genomic level in our three sets of populations by examining allele frequency shifts in polymorphisms that have known associations with pigmentation. Our analyses addressed whether patterns of melanization are associated with variation in major-effect pigmentation genes, and if allele frequency shifts in the same genes are associated with patterns across all timescales.

### Candidate pigmentation SNPs do not show enrichment for allele frequency change

We designed our genomic analyses as a targeted method to ask how known pigmentation genes vary across space and time in our populations. Previous work has demonstrated that thousands of polymorphisms exhibit clinality and seasonality in *D. melanogaster* (Bergland et al. 2014, Machado et al. 2021, Rudman et al. 2022), so we aimed to narrow the genetic variation we examined to only candidates with an established connection to pigmentation in natural populations. We focused on polymorphisms in major- and minor-effect pigmentation genes that likely represent commonly segregating variants, and we compiled our list of candidates based on pigmentation-associated SNPs identified by GWAS in both natural *D. melanogaster* populations and inbred lines from the Drosophila Genetic Reference Panel (DGRP) (Bastide et al., 2013; Dembeck et al., 2015). This list included polymorphisms in canonical genes of the biosynthetic pathway of sclerotin and melanin, as well as smaller-effect loci in genes outside of this core pathway, and it totaled 249 SNPs associated with ∼110 genes.

We first asked whether the candidate pigmentation SNPs were enriched as a group for latitudinal allele frequency variation relative to the genomic average. We tested for group enrichment of pigmentation SNPs by comparing their distribution to permutated distributions of matched control SNPs, and we looked for a significant magnitude of allele frequency change (see Methods). We determined that our candidate pigmentation SNPs were not significantly more clinal relative to matched controls (asymptotic two-sample KS test, *p* = .5233; Fig. 3; Table S2). We then repeated these analyses for our wild (*p* = .5174) and experimental (*p* = .8231) seasonal populations in the years corresponding to our phenotypic data, and similarly found that pigmentation SNPs were not enriched as a group for seasonal allele frequency change above the expectation by chance (Fig. 3; Table S2). These results were not entirely surprising, as it is possible *D. melanogaster* pigmentation patterns are being driven by selection across few loci, similar to other species (Hoekstra et al., 2006; Miller et al., 2007). Therefore, we next analyzed allele frequency shifts in each individual pigmentation SNP across all spatiotemporal scales.

**Fig 3.**
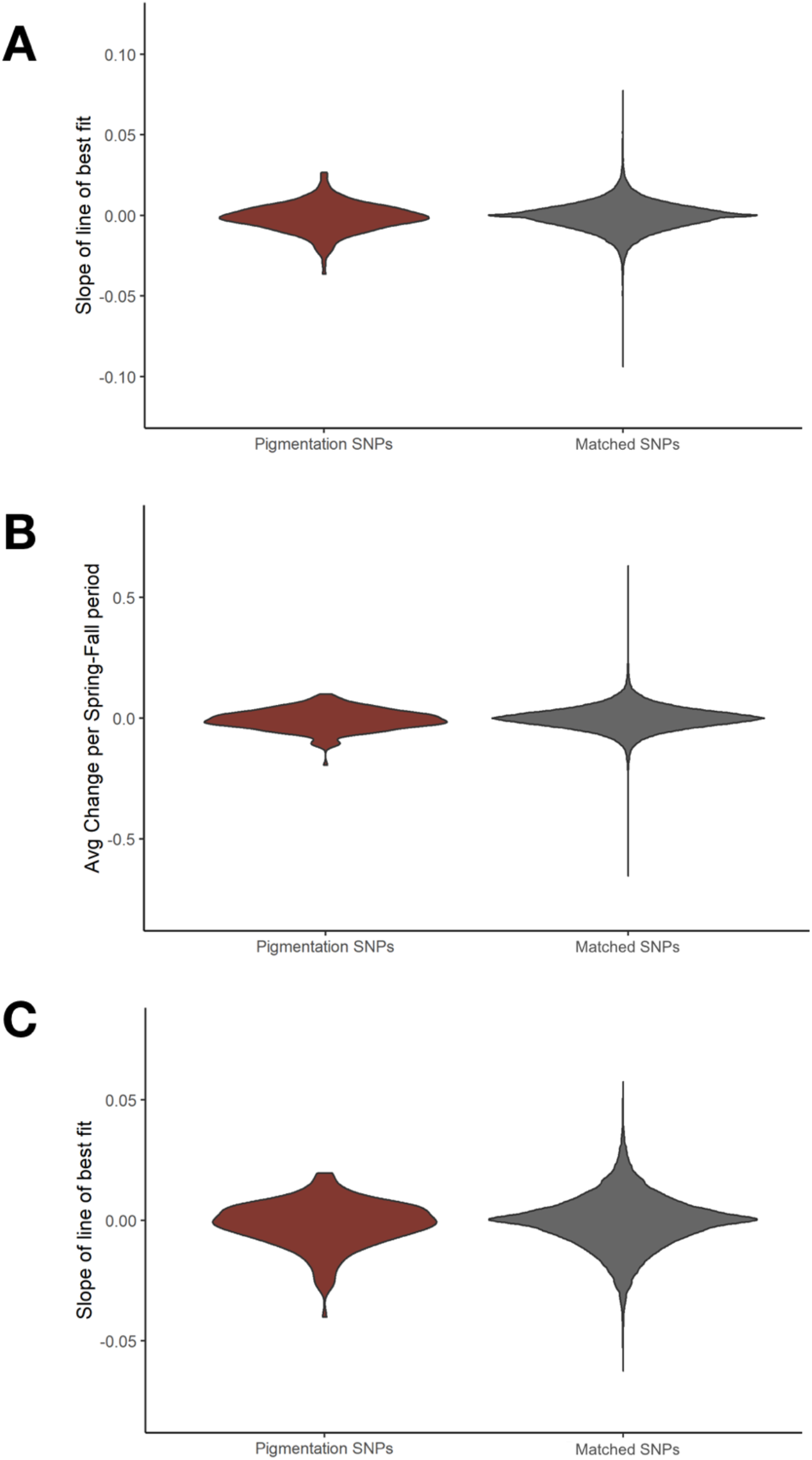
Distributions of candidate pigmentation SNPs vs. bootstrap resamples of matched SNPs over space and time. Candidate pigmentation SNPs (maroon) are not enriched as a group relative to matched controls (gray) across (A) latitudinal, (B) seasonal, or (C) experimental populations. Candidate SNPs were selected based on associations with pigmentation from GWAS of natural populations, and enrichment of candidate pigmentation SNPs was determined relative to groups of matched, control SNPs.

### Individual pigmentation SNPs show significant allele frequency shifts across latitudinal, seasonal, and experimental populations

We compiled a list of matched control SNPs for each pigmentation SNP, and we then generated a null distribution for each pigmentation SNP from the allele frequency changes of its matched SNPs. This enabled us to assess whether pigmentation SNPs exhibited significant allele frequency shifts relative to control SNPs (Table S3A-C). We first identified SNPs that exhibited latitudinal clinality, and they were located in *tan, bab1, bab2*, a *bab1* binding site, *TrpA1, dally*, and *dpr10* (Fig. 4; Fig. S1; Table S3A). We next examined seasonally evolving loci in the populations sampled from our focal orchard in Pennsylvania, cataloging a set of SNPs that (I) evolved rapidly over the course of ∼15 generations, and (II) did so in a recurrent manner across 6 years. We observed significant, repeated allele frequency shifts between early- and late-season flies at several loci, including sites within or nearby *tan*, a *bab1* binding site, *Doc1, Doc2, CG1887*, and *CG9134* (Fig. 4; Fig. S2; Table S3B). Finally, we analyzed genomic patterns in our experimental orchard. Here, we examined SNPs that exhibited parallel shifts across all replicated populations, and our experimental design eliminated demographic confounds that may be present in the wild populations. The majority of pigmentation SNPs began at a low or intermediate starting frequency (Fig. S3), and we identified a small set of loci that exhibited detectable allele frequency shifts over time. These SNPs rapidly changed in parallel across all nine mesocosms in fewer than 15 generations, and they included *tan, ZnT35C*, and possible binding sites for *inv* and *chinmo* (Fig. 4; Fig. S4; Table S3C). Altogether, our analyses uncovered significant allele frequency shifts for several SNPs in both major-(i.e., *tan, bab1, bab2*) and minor-effect pigmentation genes across all three spatiotemporal scales, but the sets of significant loci varied by context. Additionally, some of the significant SNPs in each context showed countergradient allele frequency shifts to phenotypic patterns (Fig. S1; Fig. S2; Fig. S4).

**Fig 4.**
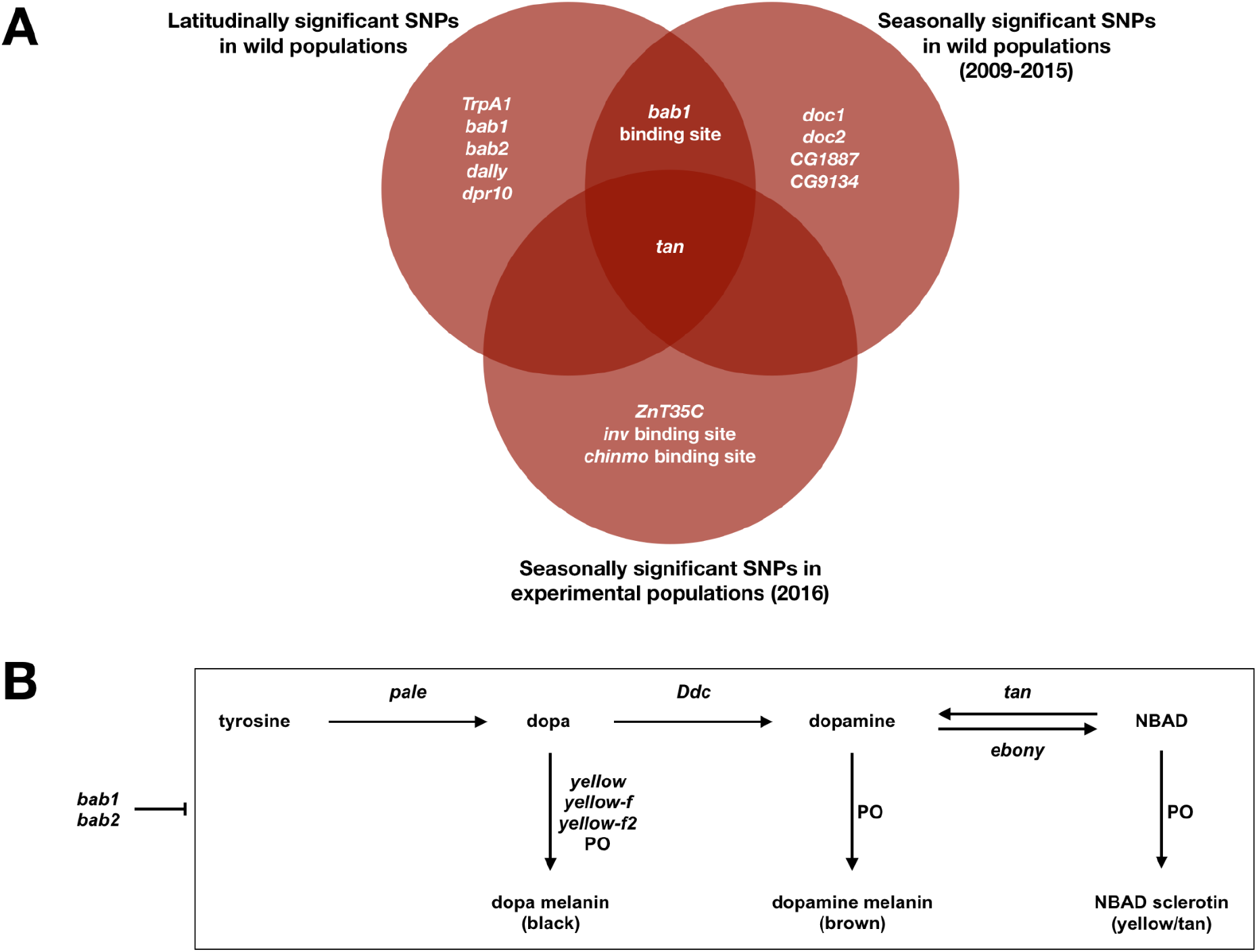
Genes harboring SNPs with significant allele frequency shifts over latitude, seasons in wild populations, and seasons in experimental populations. (A) Diagram of pigmentation genes containing significant SNPs from populations in each natural context. Significance of candidate pigmentation SNPs was determined relative to matched SNPs, and concordant genes across contexts are displayed. The directionality of allele frequency shifts was co-gradient with observed phenotypic patterns for some SNPs, while other SNPs had countergradient allele frequency shifts. (B) Simplified biosynthetic pathway displaying the canonical pigmentation genes, as well as the relationship between *bab1* and *bab2* to the pathway. *Bab* modulates the pathway by repressing *tan* and *yellow* (Salomone et al. 2013, Roeske et al. 2018), as well as *Ddc* (Gibert et al. 2007).

## Discussion

Here, we provide the first demonstration that abdominal pigmentation in *D. melanogaster* adapts rapidly and predictably as a response to differential environmental conditions, and we uncover shifts in known pigmentation genes that are associated with this change in phenotype. We first found that melanization increases with latitude in North America, mirroring latitudinal clines for pigmentation on other continents (Das, 1995; Munjal et al., 1997; Parkash et al., 2008; Rajpurohit et al., 2008a; Telonis-Scott et al., 2011). We then explored seasonal patterns in a wild population and found that flies repeatedly evolved lighter pigmentation from spring to fall over six years. Finally, we recapitulated these seasonal shifts in replicated experimental populations in the field, eliminating any effects of demography while maintaining a natural context. The shifts we observed in our experimental populations were equivalent in both magnitude and direction to the patterns we observed in wild populations, and they were established across nine replicate populations in less than fifteen generations. These data indicate that pigmentation is rapidly adapting in our experimental populations, and therefore also suggest that pigmentation is under selection in wild populations as a response to temperature or other ecological factors that co-vary with latitude and season. Thus, we found that *D. melanogaster* pigmentation demonstrated concordant and highly predictable adaptive patterns across all spatiotemporal scales.

An important caveat to our phenotypic findings is that the effects of *Drosophila* pigmentation on fitness have not been established in natural populations; it is possible that pigmentation may be under indirect selection due to correlations with other traits (Rajpurohit et al., 2016), rather than direct selection on the degree of melanization itself. Thermoregulation, UV tolerance, desiccation tolerance, cuticle strength, and wound repair have been posited as fitness benefits associated with *Drosophila* pigmentation (Freoa et al., 2023; Rajpurohit et al., 2008b; Subasi et al., 2024). However, opposite coloration patterns are observed across various species inhabiting regions with similar biotic and abiotic conditions (Kronforst et al., 2012), which obscures the adaptive significance of pigmentation phenotypes. Future experiments will examine if pigmentation is under direct selection by (I) manipulating the putative causative agents of selection (e.g., temperature) and observing consequent effects on pigmentation in our experimental orchard, and (II) testing genetic correlations of pigmentation and other fitness-related traits using a quantitative genetics framework.

Our genomic analyses revealed that common variants of major-effect pigmentation genes are associated with shifts in melanization in each spatiotemporal context, including over ecological timescales: these included *tan, bab1*, and *bab2* (Fig. 4). This finding reflects the genetic basis of pigmentation variation in natural populations across several taxa, which also experience selection at key loci for pigmentation biosynthesis (Anderson et al., 2009; Barrett et al., 2019; Gratten et al., 2007; Gratten et al., 2010; Hoekstra et al., 2006; Jones et al., 2018; Manceau et al., 2011; Miller et al., 2007; van’t Hof et al., 2016). However, while we found that canonical pigmentation genes are associated with spatiotemporal phenotypic variation in *D. melanogaster*, the regulation of melanization appears to be more nuanced. We observed that SNPs in numerous minor-effect pigmentation genes also shifted significantly across all scales, and a distinct combination of major- and minor-effect alleles were associated with melanization in each context. We also failed to identify significant patterns for alleles of other canonical genes involved in pigmentation biosynthesis, including *ebony*, which is associated with altitudinal variation for melanization (Pool & Aquadro, 2007). Thus, adaptive pigmentation patterns appear to be determined by the combined effects of many loci that are incorporated in a context-dependent fashion, rather than exclusively by canonical pigmentation genes.

While a nonoverlapping set of SNPs shifted significantly at the latitudinal, seasonal, and experimental scales, we did observe some parallelism at the gene-level: SNPs in *tan* and *bab1* shifted in multiple contexts (Fig. 4). However, the magnitude of these shifts over short timescales appears insufficient to fully account for phenotypic evolution (Fig. S2; Fig. S4). Many of the major-effect *Drosophila* pigmentation genes are highly pleiotropic (Drapeau et al., 2003; Godt et al., 1993; Jacobs, 1985; Kopp et al., 2000; Massey et al., 2019; Shaw et al., 2000; True et al., 2005; Wilson et al., 1976), so the magnitude of allele frequency shifts in canonical genes may have been limited by the degree to which pleiotropic constraints can be alleviated over each timescale (Pavličev & Cheverud, 2015). We also observed that much of the spatial and temporal allelic variation exhibited countergradient patterns, in which allele frequency differences did not match phenotypes observed (Fig. S1; Fig. S2; Fig S4). Countergradient patterns have been previously reported for genetic polymorphisms in *D. melanogaster* (Durmaz et al., 2019; Lee et al., 2013; Paaby et al., 2014; Yu & Bergland, 2022), contrasting the simple prediction that allele frequency shifts would be concordant with phenotype. Interactions among multiple combinations of alleles can produce redundant phenotypic patterns, and this can generate allele frequency shifts that conflict in directionality with phenotypic clines (Lotterhos, 2023). Additionally, epistatic interactions influence the effects of individual SNPs on quantitative trait variation across genetic backgrounds in *D. melanogaster* (Huang et al., 2012); thus, epistasis may modulate the effects of a given allele on phenotype across contexts (Noble et al., 2017; Bernstein et al., 2019). Finally, it is important to note that our genomic analyses examined known, common variation in pigmentation genes, so we may not have captured patterns of evolution in any novel, noncoding, or rare variants that contribute to phenotype (Tennessen et al., 2012).

In conclusion, our findings illustrate that *D. melanogaster* abdominal pigmentation evolves as a deterministic response to predictable shifts in environmental conditions that vary over both space and time. Yet, while strikingly parallel pigmentation patterns arose across all spatiotemporal scales, the underlying association between genotype and phenotype varied across all datasets. Therefore, we found that adaptive phenotypic patterns are produced in a predictable fashion over all timescales analyzed, but the genetic basis of complex trait evolution appears context-dependent. This study contributes to our growing understanding of how patterns of genomic and phenotypic evolution integrate in the field, and whether evolution is predictable in natural populations. Thus, our findings will better inform our understanding of the mechanisms through which populations adapt to rapid environmental change.

## Author Contributions

S.B., S.R., and P.S. designed the study. Field samples of *D. melanogaster* were collected by E.L.B., A.O.B., and P.S. The 2016 experimental orchard study was conducted by M.C. Berner, S.R., and P.S. Pigmentation data was collected by S.B., N.J.B., and S.R., and phenotypic analyses were conducted by S.B. DNA sequencing and processing of allele frequency data for the 2016 orchard experiment was completed by S.I.G. and M.C. Bitter. J.A.R. and D.A.P. designed and conducted genomic analyses. S.B. and P.S. wrote the manuscript, with feedback provided by all authors.

## Acknowledgements

We thank Heloise Bastide for contributing additional datasets associated with Bastide et al. (2013), which enabled us to assess for SNP concordance with phenotypic patterns. We thank each orchard that generously allowed us to sample their fly populations. Finally, we thank current and past members of the P.S., S.R., and D.A.P. research groups for their feedback on this manuscript.

## Data and Code Availability

Latitudinal and seasonal genomic data can be accessed via the DEST dataset, v.1 (Kapun et al., 2021). Sequencing data for the mesocosm experiment used in this study have been uploaded to the NCBI SRA under BioProject ID PRJNA1141556 and will be publicly available upon manuscript acceptance. Raw data and scripts have been uploaded to the Github repository, “skylerberardi/Papers/Berardi_2024”.

## Supplementary Materials

**Figure S1.**
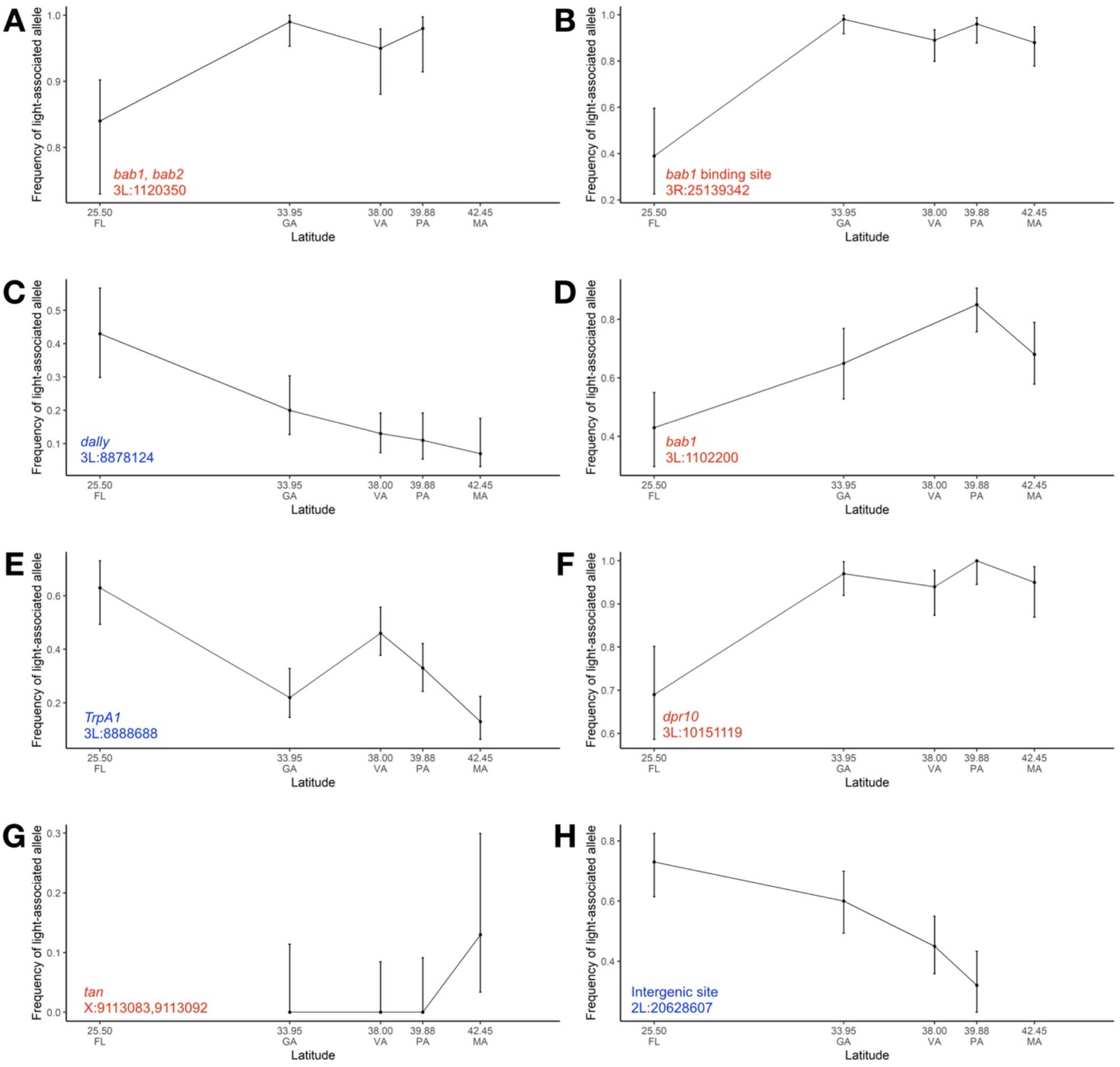
Individual pigmentation SNPs with significant allele frequency shifts across the North American latitudinal cline. Significant SNPs were identified by performing a linear regression of the allele frequencies across latitudes, and then determining if the regression slope for each candidate pigmentation SNP was in the tail of the null distribution created from the slopes of its matched SNPs. The allele frequency of the light-associated allele across latitudes is plotted to illustrate latitudinal differences in SNP frequencies, and error bars represent the 80% confidence interval. Some allele frequency clines were concordant with phenotypic clines (blue text), while other SNPs demonstrated reverse clines (red text).

**Figure S2.**
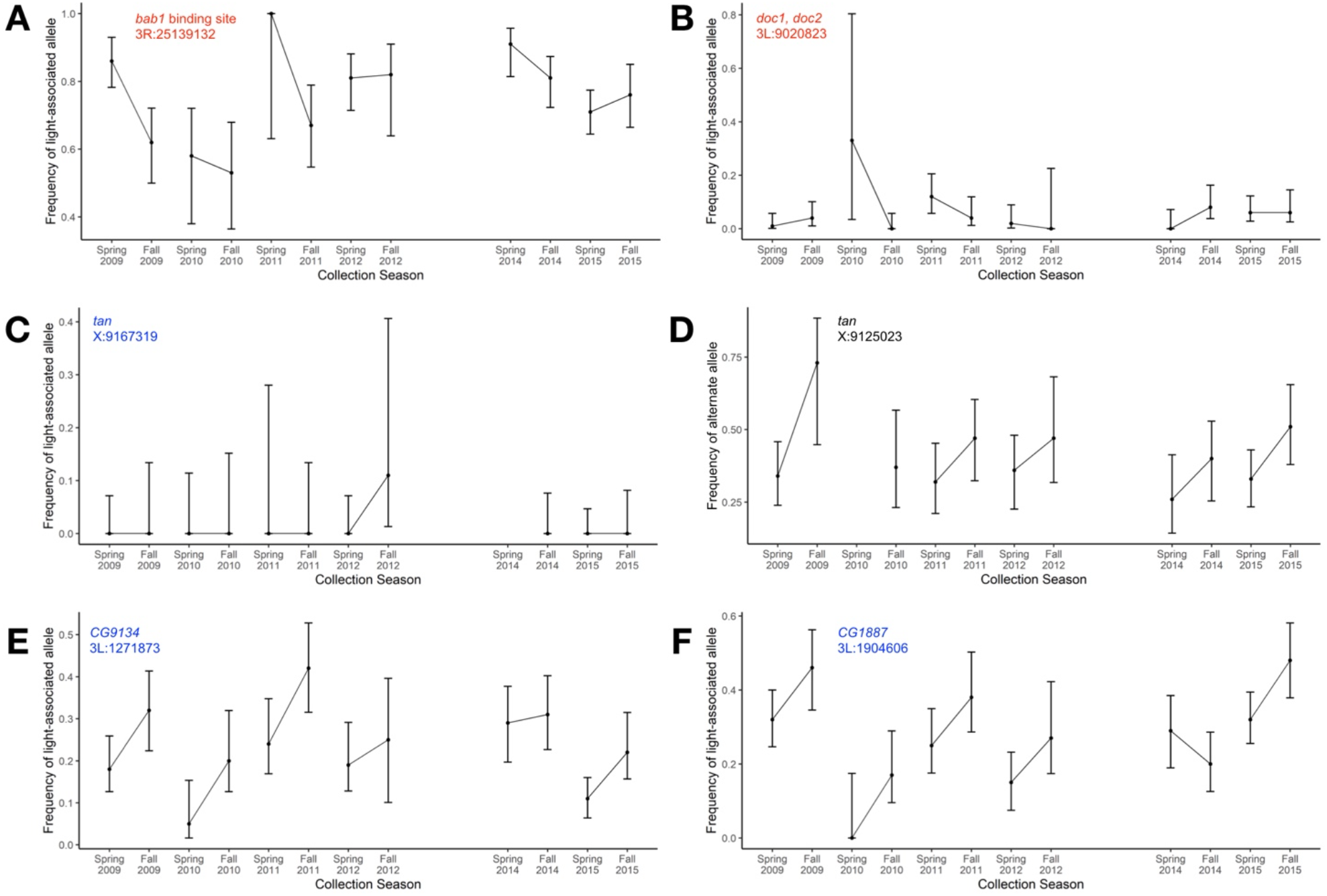
Individual pigmentation SNPs with significant allele frequency fluctuations from spring to fall across multiple years. We determined whether repeated allele frequency shifts were significant by calculating the allele frequency changes from spring to fall in each year and determining the average shift per year for each SNP. We then measured whether the shifts for each candidate pigmentation SNP were in the tail of the null distribution generated from their matched SNPs. The allele frequency of the light-associated allele is plotted across collection seasons to show parallel shifts in SNP frequencies across seasons in multiple years, and error bars represent the 80% confidence interval. Some SNPs exhibited more consistent patterns than others across years, and SNPs showed both co-gradient (blue text) and counter-gradient (red text) patterns with the phenotypic patterns observed. The directionality for one of the *tan* SNPs (X:9125023) was unspecified in the Bastide et al. (2013) dataset (black text).

**Figure S3.**
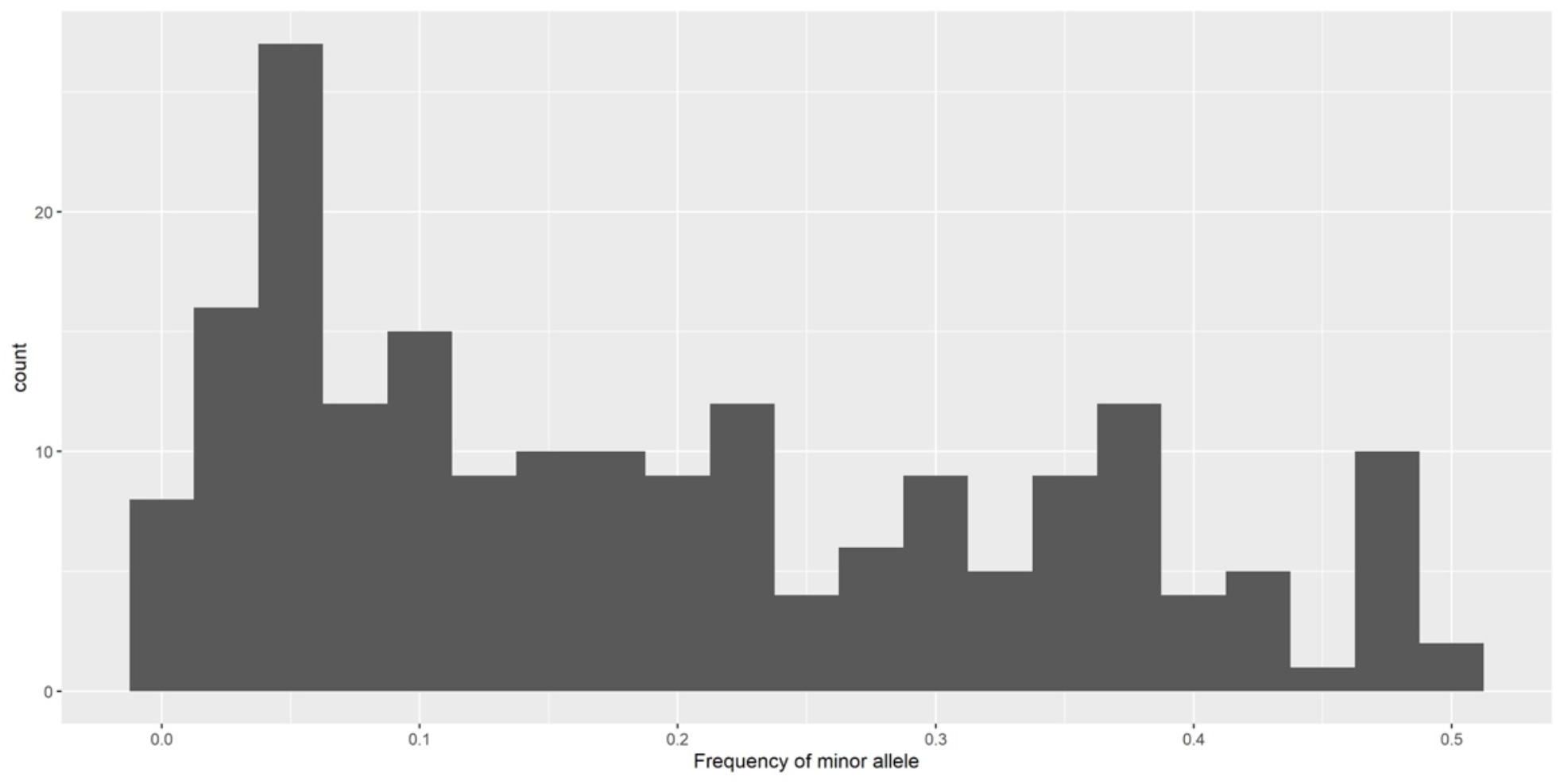
Folded site frequency spectrum displaying the starting frequency of pigmentation SNP alternate alleles in the experimental orchard. The majority of pigmentation SNPs had low or intermediate minor allele frequencies in the baseline population for the experimental orchard, with the largest proportion of initial frequencies falling between 0 and 0.1.

**Figure S4.**
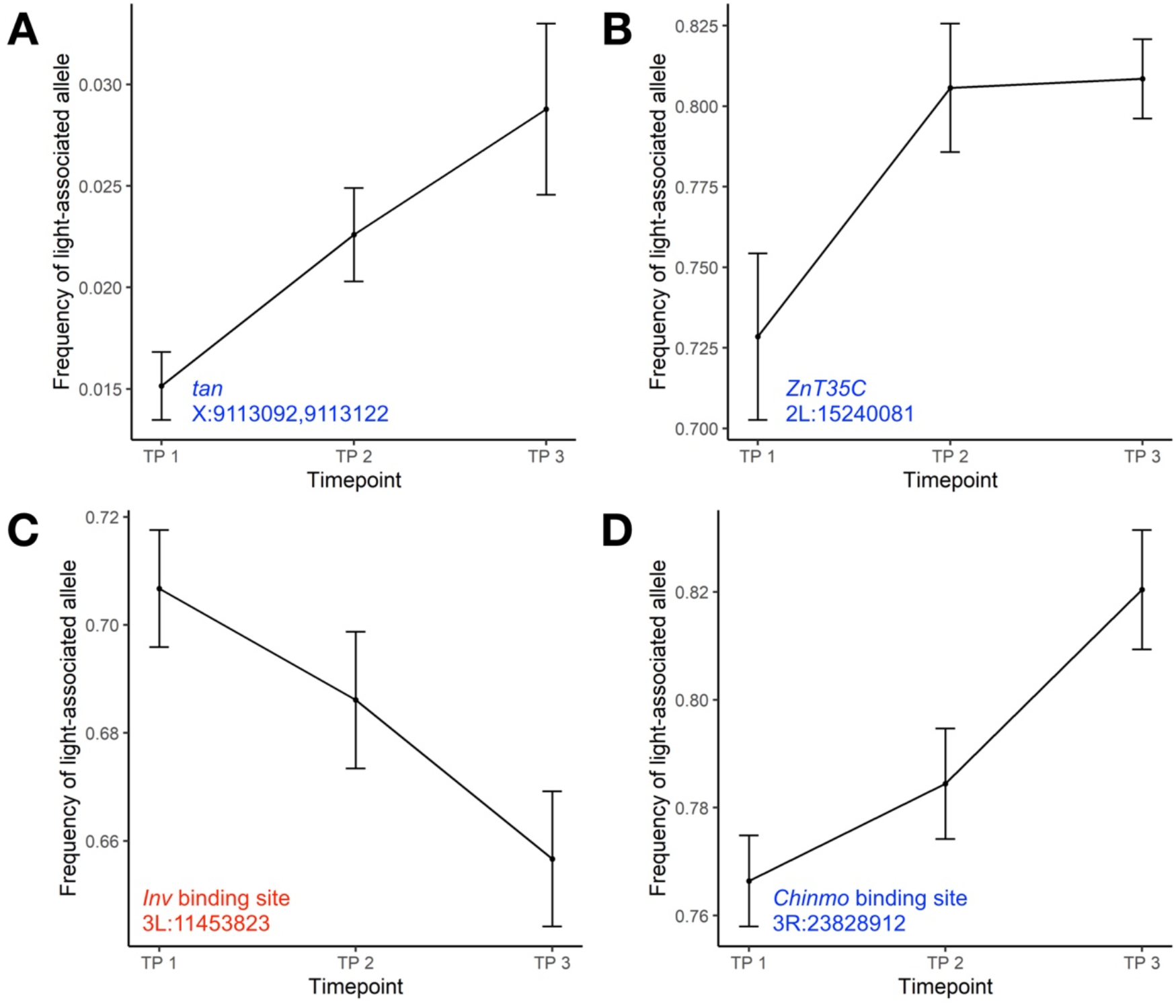
Individual pigmentation SNPs with significant allele frequency shifts across seasons in replicate experimental orchard populations. Significance was determined by first performing a linear regression of the allele frequencies across timepoints, and then identifying if the regression slope for the candidate SNP was in the tail of the null distribution created from the slopes of the matched SNPs. The allele frequency and standard error of the light-associated allele is plotted here to show the movement of SNP frequencies over time. The directionality of allele frequency shifts was concordant with observed phenotypic patterns for some SNPs (blue text), while other SNPs had countergradient allele frequency shifts (red text). (N=10 cages. TP1=Aug. 5, TP2=Sept. 5, TP3=Oct. 5).

**Table S1.** Summary table of linear mixed effects model statistics for phenotypic patterns. (A) We determined whether pigmentation varies significantly over latitudes by running a linear mixed effects model using the function lme (R package ‘nlme’, v.3.1.164). Here, latitude was the fixed effect and isofemale line was included as a random effect. (B) To examine whether pigmentation shifts significantly across seasons in the wild populations (Media, PA), we similarly ran a linear mixed effects model where year, season, and their interaction were fixed effects, and isofemale line was a random effect. (C) Finally, we probed for significant pigmentation patterns across timepoints in our experimental orchard (Philadelphia, PA) by running a linear mixed effects model where timepoint was the fixed effect and mesocosm was a random effect nested within timepoint.

|  | numDF | denDF | F-value | p-value |
| --- | --- | --- | --- | --- |
| (Intercept) | 1 | 157 | 2909.3350 | < .0001 |
| Latitude | 5 | 157 | 11.1707 | < .0001 |

**B** Experiment: seasonal wild populations Formula: Pigmentation Scores ~ Year\*Season, random=~1|Isofemale Line
|  | numDF | denDF | F-value | p-value |
| --- | --- | --- | --- | --- |
| (Intercept) | 1 | 148 | 5606.354 | < .0001 |
| Year | 5 | 148 | 13.931 | < .0001 |
| Season | 1 | 148 | 174.433 | < .0001 |
| Year:Season | 5 | 148 | 11.517 | < .0001 |

**C** Experiment: seasonal experimental populations Formula: Pigmentation Scores ~ Timepoint, random=~1|Mesocosm/Timepoint
|  | numDF | denDF | F-value | p-value |
| --- | --- | --- | --- | --- |
| (Intercept) | 1 | 8 | 2658.3068 | < .0001 |
| Timepoint | 1 | 8 | 111.2574 | < .0001 |

**Table S2.** Testing pigmentation SNP enrichment as a group. Considering all pigmentation SNPs as a group, we compared the distribution of pigmentation SNPs to the distributions of 1,000 matched control SNP groups. We assessed for enrichment of pigmentation SNPs by determining whether they exhibited a significant magnitude of absolute change across spatiotemporal gradients relative to the matched control groups. We completed these analyses using an asymptotic two-sample Kolmogorov-Smirnov test with a one-sided distribution, and we found that pigmentation SNPs as a group were not enriched in all three contexts.

|  | D<sup>-</sup> | *p-value* | Significant difference between distributions |
| --- | --- | --- | --- |
| East Coast Latitudinal Cline Populations | 0.037957 | 0.5233 | No |
| Seasonal Populations (Media, PA) | 0.038118 | 0.5174 | No |
| Seasonal Populations in Experimental Orchard (Philadelphia, PA) | 0.022646 | 0.8231 | No |

**Table S3.** Individual pigmentation SNPs with significant allele frequency shifts across each spatiotemporal gradient. (A) List of pigmentation SNPs that exhibited significant allele frequency shifts across latitudinal populations sampled along the East Coast of the U.S. (B) Pigmentation SNPs with significant shifts across seasons in a population sampled from Media, PA, over six years (2009-2012 and 2014-2015). (C) Pigmentation SNPs that shifted significantly across seasons in replicate experimental populations housed in field mesocosms (Philadelphia, PA).

| **A** | HIT | SLOPE | 10th percentile of *p*-value distribution | 50th percentile of *p*-value distribution | 90th percentile of *p*-value distribution | GENE | Concordant with phenotypic directionality? |
| --- | --- | --- | --- | --- | --- | --- | --- |
|  | 2L_20628607 | 0.026569095 | 0.01 | 0.02 | 0.04 | intergenic | yes |
|  | 3L_8888688 | -0.022554119 | 0.009 | 0.02 | 0.04 | *TrpA1* | yes |
|  | 3L_1120350 | -0.009173534 | 0.03 | 0.05 | 0.081 | *bab1, bab2* | no |
|  | 3L_8878124 | -0.021389754 | 0 | 0.01 | 0.021 | *dally* | yes |
|  | 3L_10151119 | -0.016300426 | 0 | 0.01 | 0.02 | *dpr10* | no |
|  | 3L_1102200 | -0.019337781 | 0.02 | 0.05 | 0.09 | *bab* | no |
|  | 3R_25139342 | -0.030024034 | 0 | 0.01 | 0.01 | binding site for *bab1* | no |
|  | X_9113092 | 0.011103075 | 0 | 0 | 0.02 | *tan* | no |
|  | X_9113083 | 0.013121816 | 0 | 0 | 0.02 | *tan* | no |
| **B** | HIT | SEASON METRIC | 10th percentile of *p*-value distribution | 50th percentile of *p*-value distribution | 90th percentile of *p*-value distribution | GENE | Concordant with phenotypic directionality? |
|  | 3L_9020823 | -0.053333333 | 0.02 | 0.045 | 0.07 | *Doc1, Doc2* | no |
|  | 3L_1904606 | 0.105 | 0.02 | 0.04 | 0.07 | *CG1887* | yes |
|  | 3L_1271873 | 0.11 | 0 | 0.02 | 0.04 | *CG9134* | yes |
|  | X_9167319 | 0.022 | 0.02 | 0.04 | 0.08 | *tan* | yes |
|  | X_9125023 | 0.194 | 0 | 0.01 | 0.02 | *tan* | SNP association with phenotype not given |
|  | 3R_25139132 | 0.11 | 0.029 | 0.04 | 0.06 | binding site for *bab1* | no |
| **C** | HIT | SLOPE | 10th percentile of *p*-value distribution | 50th percentile of *p*-value distribution | 90th percentile of *p*-value distribution | GENE | Concordant with phenotypic directionality? |
|  | 2L_15240081 | -0.040018867 | 0 | 0 | 0.011 | *ZnT35C*, binding site for *HSA, sbb* | yes |
|  | 3L_11453823 | -0.025027783 | 0 | 0.02 | 0.03 | possible binding site for *inv* | no |
|  | X_9113122 | 0.006818617 | 0.01 | 0.03 | 0.05 | *tan* | yes |
|  | X_9113092 | 0.006818617 | 0.01 | 0.04 | 0.061 | *tan* | yes |
|  | 3R_23828912 | -0.027007122 | 0.03 | 0.05 | 0.08 | possible binding site for *chinmo* | yes |

## References

Allen, J. A. (1877). The influence of physical conditions in the genesis of species. Radical Review, 1, 108–140.

Anderson, T. M., vonHoldt, B. M., Candille, S. I., Musiani, M., Greco, C., Stahler, D. R., Smith, D. W., Padhukasahasram, B., Randi, E., Leonard, J. A., Bustamante, C. D., Ostrander, E. A., Tang, H., Wayne, R. K., & Barsh, G. S. (2009). Molecular and evolutionary history of melanism in North American gray wolves. Science, 323(5919), 1339–1343. 10.1126/science.1165448

Barrett, R. D., Rogers, S. M., & Schluter, D. (2008). Natural selection on a major armor gene in threespine stickleback. Science, 322(5899), 255–257. 10.1126/science.1159978

Barrett, R. D., Laurent, S., Mallarino, R., Pfeifer, S. P., Xu, C. C., Foll, M., Wakamatsu, K., Duke-Cohan, J. S., Jensen, J. D., & Hoekstra, H. E. (2019). Linking a mutation to survival in wild mice. Science, 363(6426), 499–504. 10.1126/science.aav3824

Bastide, H., Betancourt, A., Nolte, V., Tobler, R., Stöbe, P., Futschik, A., & Schlötterer, C. (2013). A genome-wide, fine-scale map of natural pigmentation variation in Drosophila melanogaster. PLoS Genetics, 9(6), e1003534. 10.1371/journal.pgen.1003534

Bastide, H., Yassin, A., Johanning, E. J., & Pool, J. E. (2014). Pigmentation in Drosophila melanogaster reaches its maximum in Ethiopia and correlates most strongly with ultra-violet radiation in sub-Saharan Africa. BMC Evolutionary Biology, 14, 1–14. 10.1186/s12862-014-0179-y

Behrman, E. L., Watson, S. S., O’Brien, K. R., Heschel, M. S., & Schmidt, P. S. (2015). Seasonal variation in life history traits in two Drosophila species. Journal of Evolutionary Biology, 28(9), 1691–1704. 10.1111/jeb.12690

Behrman, E. L., Howick, V. M., Kapun, M., Staubach, F., Bergland, A. O., Petrov, D. A., Lazzaro, B., & Schmidt, P. S. (2018). Rapid seasonal evolution in innate immunity of wild Drosophila melanogaster. Proceedings of the Royal Society B: Biological Sciences, 285(1870), 20172599. 10.1098/rspb.2017.2599

Behrman, E. L., & Schmidt, P. (2022). How predictable is rapid evolution? BioRxiv, 2022–10. 10.1101/2022.10.27.514123

Bergland, A. O., Behrman, E. L., O’Brien, K. R., Schmidt, P. S., & Petrov, D. A. (2014). Genomic evidence of rapid and stable adaptive oscillations over seasonal time scales in Drosophila. PLoS Genetics, 10(11), e1004775. 10.1371/journal.pgen.1004775

Bergland, A. O., Tobler, R., González, J., Schmidt, P., & Petrov, D. (2016). Secondary contact and local adaptation contribute to genome-wide patterns of clinal variation in Drosophila melanogaster. Molecular Ecology, 25(5), 1157–1174. 10.1111/mec.13455

Bernstein, M. R., Zdraljevic, S., Andersen, E. C., & Rockman, M. V. (2019). Tightly linked antagonistic-effect loci underlie polygenic phenotypic variation in C. elegans. Evolution Letters, 3(5), 462–473. 10.1002/evl3.139

Bitter, M. C., Berardi, S., Oken, H., Huynh, A., Lappo, E., Schmidt, P., & Petrov, D. A. (2024). Continuously fluctuating selection reveals fine granularity of adaptation. Nature, 634(8033), 389–396. 10.1038/s41586-024-07834-x

Couderc, J. L., Godt, D., Zollman, S., Chen, J., Li, M., Tiong, S., Cramton, S. E., Sahut-Barnola, I., & Laski, F. A. (2002). The bric à brac locus consists of two paralogous genes encoding BTB/POZ domain proteins and acts as a homeotic and morphogenetic regulator of imaginal development in Drosophila. Development, 129(10), 2419–2433. 10.1242/dev.129.10.2419

Das, A. (1995). Abdominal pigmentation in Drosophila melanogaster females from natural Indian populations. Journal of Zoological Systematics and Evolutionary Research, 33(2), 84–87. 10.1111/j.1439-0469.1995.tb00213.x

David, J. R., Capy, P., Payant, V., & Tsakas, S. (1985). Thoracic trident pigmentation in Drosophila melanogaster: differentiation of geographical populations. Génétique Sélection Évolution, 17(2), 211–224.

David, J. R., Capy, P., & Gauthier, J. P. (1990). Abdominal pigmentation and growth temperature in Drosophila melanogaster: similarities and differences in the norms of reaction of successive segments. Journal of Evolutionary Biology, 3(5-6), 429–445. 10.1046/j.1420-9101.1990.3050429.x

Dembeck, L. M., Huang, W., Magwire, M. M., Lawrence, F., Lyman, R. F., & Mackay, T. F. (2015). Genetic architecture of abdominal pigmentation in Drosophila melanogaster. PLoS Genetics, 11(5), e1005163. 10.1371/journal.pgen.1005163

Dev, K., Chahal, J., Parkash, R., & Kataria, S. K. (2013). Correlated changes in body melanisation and mating traits of Drosophila melanogaster: A seasonal analysis. Evolutionary Biology, 40, 366–376. 10.1007/s11692-012-9220-5

Dobzhansky, T. (1943). Genetics of natural populations IX. Temporal changes in the composition of populations of Drosophila pseudoobscura. Genetics, 28(2), 162. 10.1093/genetics/28.2.162

dos Santos, G., Schroeder, A. J., Goodman, J. L., Strelets, V. B., Crosby, M. A., Thurmond, J., Emmert, D. B., Gelbart, W. M., & FlyBase Consortium. (2015). FlyBase: introduction of the Drosophila melanogaster Release 6 reference genome assembly and large-scale migration of genome annotations. Nucleic Acids Research, 43(D1), D690–D697. 10.1093/nar/gku1099

Drapeau, M. D., Radovic, A., Wittkopp, P. J., & Long, A. D. (2003). A gene necessary for normal male courtship, yellow, acts downstream of fruitless in the Drosophila melanogaster larval brain. Journal of Neurobiology, 55(1), 53–72. 10.1002/neu.10196

Durmaz, E., Rajpurohit, S., Betancourt, N., Fabian, D. K., Kapun, M., Schmidt, P., & Flatt, T. (2019). A clinal polymorphism in the insulin signaling transcription factor foxo contributes to life-history adaptation in Drosophila. Evolution, 73(9), 1774–1792. 10.1111/evo.13759

Enbody, E. D., Sendell-Price, A. T., Sprehn, C. G., Rubin, C. J., Visscher, P. M., Grant, B. R., Grant, P. R., & Andersson, L. (2023). Community-wide genome sequencing reveals 30 years of Darwin’s finch evolution. Science, 381(6665), eadf6218. 10.1126/science.adf6218

Endler, J. A. (1977). Geographic variation, speciation, and clines (No. 10). Princeton University Press.

Endler, L., Betancourt, A. J., Nolte, V., & Schlötterer, C. (2016). Reconciling differences in pool-GWAS between populations: a case study of female abdominal pigmentation in Drosophila melanogaster. Genetics, 202(2), 843–855. 10.1534/genetics.115.183376

Erickson, P. A., Weller, C. A., Song, D. Y., Bangerter, A. S., Schmidt, P., & Bergland, A. O. (2020). Unique genetic signatures of local adaptation over space and time for diapause, an ecologically relevant complex trait, in Drosophila melanogaster. PLoS Genetics, 16(11), e1009110. 10.1371/journal.pgen.1009110

Freoa, L., Chevin, L. M., Christol, P., Méléard, S., Rera, M., Véber, A., & Gibert, J. M. (2023). Drosophilids with darker cuticle have higher body temperature under light. Scientific Reports, 13(1), 3513. 10.1038/s41598-023-30652-6

Gibert, J. M., Peronnet, F., & Schlötterer, C. (2007). Phenotypic plasticity in Drosophila pigmentation caused by temperature sensitivity of a chromatin regulator network. PLoS Genetics, 3(2), e30. 10.1371/journal.pgen.0030030

Godt, D., Couderc, J. L., Cramton, S. E., & Laski, F. A. (1993). Pattern formation in the limbs of Drosophila: bric à brac is expressed in both a gradient and a wave-like pattern and is required for specification and proper segmentation of the tarsus. Development, 119(3), 799–812. 10.1242/dev.119.3.799

Grainger, T. N., Rudman, S. M., Schmidt, P., & Levine, J. M. (2021). Competitive history shapes rapid evolution in a seasonal climate. Proceedings of the National Academy of Sciences, 118(6), e2015772118. 10.1073/pnas.2015772118

Gratten, J., Beraldi, D., Lowder, B. V., McRae, A. F., Visscher, P. M., Pemberton, J. M., & Slate, J. (2007). Compelling evidence that a single nucleotide substitution in TYRP1 is responsible for coat-colour polymorphism in a free-living population of Soay sheep. Proceedings of the Royal Society B: Biological Sciences, 274(1610), 619–626. 10.1098/rspb.2006.3762

Gratten, J., Pilkington, J. G., Brown, E. A., Beraldi, D., Pemberton, J. M., & Slate, J. (2010). The genetic basis of recessive self-colour pattern in a wild sheep population. Heredity, 104(2), 206–214. 10.1038/hdy.2009.105

Hoekstra, H. E., Hirschmann, R. J., Bundey, R. A., Insel, P. A., & Crossland, J. P. (2006). A single amino acid mutation contributes to adaptive beach mouse color pattern. Science 313(5783), 101–104. 10.1126/science.1126121

Huang, W., Richards, S., Carbone, M. A., Zhu, D., Anholt, R. R., Ayroles, J. F., … & Mackay, T. F. (2012). Epistasis dominates the genetic architecture of Drosophila quantitative traits. Proceedings of the National Academy of Sciences, 109(39), 15553–15559. 10.1073/pnas.1213423109

Jacobs, M. E. (1985). Role of beta-alanine in cuticular tanning, sclerotization, and temperature regulation in Drosophila melanogaster. Journal of Insect Physiology, 31(6), 509–515. 10.1016/0022-1910(85)90099-X

Jones, M. R., Mills, L. S., Alves, P. C., Callahan, C. M., Alves, J. M., Lafferty, D. J., Jiggins, F. M., Jensen, J. D., Melo-Ferreira, J., & Good, J. M. (2018). Adaptive introgression underlies polymorphic seasonal camouflage in snowshoe hares. Science, 360(6395), 1355–1358. 10.1126/science.aar5273

Kapun, M., Nunez, J. C., Bogaerts-Márquez, M., Murga-Moreno, J., Paris, M., Outten, J., … & Bergland, A. O. (2021). Drosophila evolution over space and time (DEST): a new population genomics resource. Molecular Biology and Evolution, 38(12), 5782–5805. 10.1093/molbev/msab259

Knoll, A. H., & Carroll, S. B. (1999). Early animal evolution: emerging views from comparative biology and geology. Science, 284(5423), 2129–2137. 10.1126/science.284.5423.2129

Kopp, A., Duncan, I., & Carroll, S. B. (2000). Genetic control and evolution of sexually dimorphic characters in Drosophila. Nature, 408(6812), 553–559. 10.1038/35046017

Kopp, A., Graze, R. M., Xu, S., Carroll, S. B., & Nuzhdin, S. V. (2003). Quantitative trait loci responsible for variation in sexually dimorphic traits in Drosophila melanogaster. Genetics, 163(2), 771–787. 10.1093/genetics/163.2.771

Kostyun, J. L., Gibson, M. J., King, C. M., & Moyle, L. C. (2019). A simple genetic architecture and low constraint allow rapid floral evolution in a diverse and recently radiating plant genus. New Phytologist, 223(2), 1009–1022. 10.1111/nph.15844

Kronforst, M. R., Barsh, G. S., Kopp, A., Mallet, J., Monteiro, A., Mullen, S. P., Protas, M., Rosenblum, E. B., Schneider, C. J., & Hoekstra, H. E. (2012). Unraveling the thread of nature’s tapestry: the genetics of diversity and convergence in animal pigmentation. Pigment Cell & Melanoma Research, 25(4), 411–433. 10.1111/j.1755-148X.2012.01014.x

Lee, S. F., Eyre-Walker, Y. C., Rane, R. V., Reuter, C., Vinti, G., Rako, L., Partridge, L., & Hoffmann, A. A. (2013). Polymorphism in the neurofibromin gene, Nf1, is associated with antagonistic selection on wing size and development time in Drosophila melanogaster. Molecular Ecology, 22(10), 2716–2725. 10.1111/mec.12301

Lenski, R. E., Rose, M. R., Simpson, S. C., & Tadler, S. C. (1991). Long-term experimental evolution in Escherichia coli. I. Adaptation and divergence during 2,000 generations. The American Naturalist, 138(6), 1315–1341. 10.1086/285289

Lotterhos, K. E. (2023). The paradox of adaptive trait clines with nonclinal patterns in the underlying genes. Proceedings of the National Academy of Sciences, 120(12), e2220313120. 10.1073/pnas.2220313120

Machado, H. E., Bergland, A. O., Taylor, R., Tilk, S., Behrman, E., Dyer, K., … & Petrov, D. A. (2021). Broad geographic sampling reveals the shared basis and environmental correlates of seasonal adaptation in Drosophila. Elife, 10, e67577. 10.7554/eLife.67577

Manceau, M., Domingues, V. S., Mallarino, R., & Hoekstra, H. E. (2011). The developmental role of Agouti in color pattern evolution. Science, 331(6020), 1062–1065. 10.1126/science.1200684

Massey, J. H., & Wittkopp, P. J. (2016). The genetic basis of pigmentation differences within and between Drosophila species. Current Topics in Developmental Biology, 119, 27–61. 10.1016/bs.ctdb.2016.03.004

Massey, J. H., Akiyama, N., Bien, T., Dreisewerd, K., Wittkopp, P. J., Yew, J. Y., & Takahashi, A. (2019). Pleiotropic effects of ebony and tan on pigmentation and cuticular hydrocarbon composition in Drosophila melanogaster. Frontiers in Physiology, 10, 518. 10.3389/fphys.2019.00518

Miller, C. T., Beleza, S., Pollen, A. A., Schluter, D., Kittles, R. A., Shriver, M. D., & Kingsley, D. M. (2007). cis-Regulatory changes in Kit ligand expression and parallel evolution of pigmentation in sticklebacks and humans. Cell, 131(6), 1179–1189. 10.1016/j.cell.2007.10.055

Munjal, A. K., Karan, D., Gibert, P., Moreteau, B., Parkash, R., & David, J. R. (1997). Thoracic trident pigmentation in Drosophila melanogaster: latitudinal and altitudinal clines in Indian populations. Genetics Selection Evolution, 29(5), 601–610.

Noble, L. M., Chelo, I., Guzella, T., Afonso, B., Riccardi, D. D., Ammerman, P., … & Teotónio, H. (2017). Polygenicity and epistasis underlie fitness-proximal traits in the Caenorhabditis elegans multiparental experimental evolution (CeMEE) panel. Genetics, 207(4), 1663–1685. 10.1534/genetics.117.300406

Nosil, P., Villoutreix, R., de Carvalho, C. F., Farkas, T. E., Soria-Carrasco, V., Feder, J. L., Crespi, B. J., & Gompert, Z. (2018). Natural selection and the predictability of evolution in Timema stick insects. Science, 359(6377), 765–770. 10.1126/science.aap9125

Nunez, J. C., Lenhart, B. A., Bangerter, A., Murray, C. S., Mazzeo, G. R., Yu, Y., Nystrom, T. L., Tern, C., Erickson, P. A., & Bergland, A. O. (2024). A cosmopolitan inversion facilitates seasonal adaptation in overwintering Drosophila. Genetics, 226(2), iyad207. 10.1093/genetics/iyad207

Paaby, A. B., & Rockman, M. V. (2013). The many faces of pleiotropy. Trends in Genetics, 29(2), 66–73. 10.1016/j.tig.2012.10.010

Paaby, A. B., Bergland, A. O., Behrman, E. L., & Schmidt, P. S. (2014). A highly pleiotropic amino acid polymorphism in the Drosophila insulin receptor contributes to life-history adaptation. Evolution, 68(12), 3395–3409. 10.1111/evo.12546

Parkash, R., Rajpurohit, S., & Ramniwas, S. (2008). Changes in body melanisation and desiccation resistance in highland vs. lowland populations of D. melanogaster. Journal of Insect Physiology, 54(6), 1050–1056. 10.1016/j.jinsphys.2008.04.008

Pavličev, M., & Cheverud, J. M. (2015). Constraints evolve: context dependency of gene effects allows evolution of pleiotropy. Annual Review of Ecology, Evolution, and Systematics, 46(1), 413–434. 10.1146/annurev-ecolsys-120213-091721

Pool, J. E., & Aquadro, C. F. (2007). The genetic basis of adaptive pigmentation variation in Drosophila melanogaster. Molecular Ecology, 16(14), 2844–2851. 10.1111/j.1365-294X.2007.03324.x

Rajpurohit, S., Parkash, R., Ramniwas, S., & Singh, S. (2008a). Variations in body melanisation, ovariole number and fecundity in highland and lowland populations of Drosophila melanogaster from the Indian subcontinent. Insect Science, 15(6), 553–561. 10.1111/j.1744-7917.2008.00245.x

Rajpurohit, S., Parkash, R., & Ramniwas, S. (2008b). Body melanization and its adaptive role in thermoregulation and tolerance against desiccating conditions in drosophilids. Entomological Research, 38(1), 49–60. 10.1111/j.1748-5967.2008.00129.x

Rajpurohit, S., Richardson, R., Dean, J., Vazquez, R., Wong, G., & Schmidt, P. S. (2016). Pigmentation and fitness trade-offs through the lens of artificial selection. Biology Letters, 12(10), 20160625. 10.1098/rsbl.2016.0625

Rajpurohit, S., & Schmidt, P. S. (2016). Measuring thermal behavior in smaller insects: A case study in Drosophila melanogaster demonstrates effects of sex, geographic origin, and rearing temperature on adult behavior. Fly, 10(4), 149–161. 10.1080/19336934.2016.1194145

Rajpurohit, S., Hanus, R., Vrkoslav, V., Behrman, E. L., Bergland, A. O., Petrov, D., Cvačka, J., & Schmidt, P. S. (2017). Adaptive dynamics of cuticular hydrocarbons in Drosophila. Journal of Evolutionary Biology, 30(1), 66–80. 10.1111/jeb.12988

Rajpurohit, S., Gefen, E., Bergland, A. O., Petrov, D. A., Gibbs, A. G., & Schmidt, P. S. (2018). Spatiotemporal dynamics and genome-wide association analysis of desiccation tolerance in Drosophila melanogaster. Molecular Ecology, 27(17), 3525–3540. 10.1111/mec.14814

Roberts Kingman, G. A., Vyas, D. N., Jones, F. C., Brady, S. D., Chen, H. I., Reid, K., … & Veeramah, K. R. (2021). Predicting future from past: The genomic basis of recurrent and rapid stickleback evolution. Science Advances, 7(25), eabg5285. 10.1126/sciadv.abg5285

Roeske, M. J., Camino, E. M., Grover, S., Rebeiz, M., & Williams, T. M. (2018). Cis-regulatory evolution integrated the Bric-à-brac transcription factors into a novel fruit fly gene regulatory network. Elife, 7, e32273. 10.7554/eLife.32273

Rudman, S. M., Greenblum, S., Hughes, R. C., Rajpurohit, S., Kiratli, O., Lowder, D. B., … & Schmidt, P. (2019). Microbiome composition shapes rapid genomic adaptation of Drosophila melanogaster. Proceedings of the National Academy of Sciences, 116(40), 20025–20032. 10.1073/pnas.1907787116

Rudman, S. M., Greenblum, S. I., Rajpurohit, S., Betancourt, N. J., Hanna, J., Tilk, S., Yokoyama, T., Petrov, D. A., & Schmidt, P. (2022). Direct observation of adaptive tracking on ecological time scales in Drosophila. Science, 375(6586), eabj7484. 10.1126/science.abj7484

Salomone, J. R., Rogers, W. A., Rebeiz, M., & Williams, T. M. (2013). The evolution of Bab paralog expression and abdominal pigmentation among Sophophora fruit fly species. Evolution & Development, 15(6), 442–457. 10.1111/ede.12053

Schmidt, P. S., & Conde, D. R. (2006). Environmental heterogeneity and the maintenance of genetic variation for reproductive diapause in Drosophila melanogaster. Evolution, 60(8), 1602–1611. 10.1111/j.0014-3820.2006.tb00505.x

Shaw, P. J., Cirelli, C., Greenspan, R. J., & Tononi, G. (2000). Correlates of sleep and waking in Drosophila melanogaster. Science, 287(5459), 1834–1837. 10.1126/science.287.5459.1834

Subasi, B. S., Grabe, V., Kaltenpoth, M., Rolff, J., & Armitage, S. A. (2024). How frequently are insects wounded in the wild? A case study using Drosophila melanogaster. Royal Society Open Science, 11(6), 240256. 10.1098/rsos.240256

Telonis-Scott, M., Hoffmann, A. A., & Sgrò, C. M. (2011). The molecular genetics of clinal variation: a case study of ebony and thoracic trident pigmentation in Drosophila melanogaster from eastern Australia. Molecular Ecology, 20(10), 2100–2110. 10.1111/j.1365-294X.2011.05089.x\

Tennessen, J. A., Bigham, A. W., O’connor, T. D., Fu, W., Kenny, E. E., Gravel, S., … & NHLBI Exome Sequencing Project. (2012). Evolution and functional impact of rare coding variation from deep sequencing of human exomes. Science, 337(6090), 64–69. 10.1126/science.1219240

True, J. R. (2003). Insect melanism: the molecules matter. Trends in Ecology & Evolution, 18(12), 640–647. 10.1016/j.tree.2003.09.006

True, J. R., Yeh, S. D., Hovemann, B. T., Kemme, T., Meinertzhagen, I. A., Edwards, T. N., Liou, S. R., Han, Q. & Li, J. (2005). Drosophila tan encodes a novel hydrolase required in pigmentation and vision. PLoS Genetics, 1(5), e63. 10.1371/journal.pgen.0010063

van’t Hof, A. E., Campagne, P., Rigden, D. J., Yung, C. J., Lingley, J., Quail, M. A., Hall, N., Darby, A. C., & Saccheri, I. J. (2016). The industrial melanism mutation in British peppered moths is a transposable element. Nature, 534(7605), 102–105. 10.1038/nature17951

Wilson, R., Burnet, B., Eastwood, L., & Connolly, K. (1976). Behavioural pleiotropy of the yellow gene in Drosophila melanogaster. Genetics Research, 28(1), 75–88. 10.1017/S0016672300016748

Wittkopp, P. J., True, J. R., & Carroll, S. B. (2002). Reciprocal functions of the Drosophila Yellow and Ebony proteins in the development and evolution of pigment patterns. Development, 129(8), 1849–1858. 10.1242/dev.129.8.1849

Wittkopp, P. J., Carroll, S. B., & Kopp, A. (2003). Evolution in black and white: genetic control of pigment patterns in Drosophila. Trends in Genetics, 19(9), 495–504. 10.1016/S0168-9525(03)00194-X

Wittkopp, P. J., & Beldade, P. (2009, February). Development and evolution of insect pigmentation: genetic mechanisms and the potential consequences of pleiotropy. In Seminars in Cell & Developmental Biology (Vol. 20, No. 1, pp. 65–71). Academic Press. 10.1016/j.semcdb.2008.10.002

Wright, T. R. (1987). The genetics of biogenic amine metabolism, sclerotization, and melanization in Drosophila melanogaster. Advances in Genetics, 24, 127–222. 10.1016/S0065-2660(08)60008-5

Yu, Y., & Bergland, A. O. (2022). Distinct signals of clinal and seasonal allele frequency change at eQTLs in Drosophila melanogaster. Evolution, 76(11), 2758–2768. 10.1111/evo.14617

